# Genome-Wide Disease Screening in Early Human Embryos with Primary Template-Directed Amplification

**DOI:** 10.1101/2021.07.06.451077

**Authors:** Yuntao Xia, Veronica Gonzales-Pena, David J Klein, Joe J Luquette, Liezl Puzon, Noor Siddiqui, Vikrant Reddy, Peter Park, Barry R Behr, Charles Gawad

## Abstract

Current preimplantation genetic testing (PGT) enables the selection of embryos based on fetal aneuploidy or the presence a small number of preselected disease-associated variants. Here we present a new approach that takes advantage of the improved genome coverage and uniformity of primary template-directed amplification (PTA) to call most early embryo genetic variants accurately and reproducibly from a preimplantation biopsy. With this approach, we identified clonal and mosaic chromosomal aneuploidy, *de novo* mitochondrial variants, and variants predicted to cause mendelian and non-mendelian diseases. In addition, we utilized the genome-wide information to compute polygenic risk scores for common diseases. Although numerous computational, interpretive, and ethical challenges remain, this approach establishes the technical feasibility of screening for and preventing numerous debilitating inherited diseases.

## Introduction

The development of high-throughput sequencing technologies has enabled the rapid acceleration of our understanding of how specific genetic variants contribute to inherited diseases^1^. In addition, the creation of polygenic risk scores has provided us with new tools to assess risk of developing multifactorial diseases^2^. Still, although we can now accurately diagnose and assess risk for numerous genetic disorders, treatment options remain limited for many diseases.

Preimplantation genetic testing (PGT) has been developed to screen embryos for aneuploidy which has significantly improved implantation and subsequent pregnancy success rates^3^. In addition, strategies have been developed for identifying known genetic variants in families by first screening the parents. Clinical labs then typically use targeted PCR-based strategies to test the embryo for those known pathogenic variants^4^. However, an accurate genome-wide method for screening embryos does not currently exist and most strategies look for either aneuploidy or small genomic changes, but not both in the same embryo. This is due to the lack of genome coverage and/or uniformity of existing whole genome amplification (WGA) methods, which are required to produce sufficient quantities of DNA to sequence a few cells from a trophoectoderm (TE) biopsy.

We recently developed a much more accurate WGA method, primary template-direct amplification (PTA), which captures almost the entire genome of minute quantities of nucleic acid in a more accurate and uniform manner, enabling much more sensitive genome-wide variant calling^5^. Here we utilized PTA to create a comprehensive PGT strategy of TE biopsies that sensitively and precisely detects aneuploidy, small genomic variants, and heteroplasmy from the same embryo, allowing us to detect inherited and *de novo* genetic variation known to cause disease, as well as produce polygenic risk scores for common multifactorial diseases. Together, this approach established an approach for preventing many genetic diseases through more comprehensive screening of embryos prior to implantation.

## Results

To assess the accuracy of our PGT strategy, we performed PTA on 8 TE biopsies harvested from 4 sibling embryos that had been donated and banked for research use. As seen in Fig. 1A, two biopsies were sampled from the same embryo to determine the consistency of variant calls. Parallel analyses were performed to call chromosome ploidy, capture genomic variants, and detect heteroplasmy. In addition, we were able to conduct genome-wide variant screening for pathological variants at 7.5 million known SNP locations to produce polygenic risk scores for eleven common diseases.

**Figure 1.**
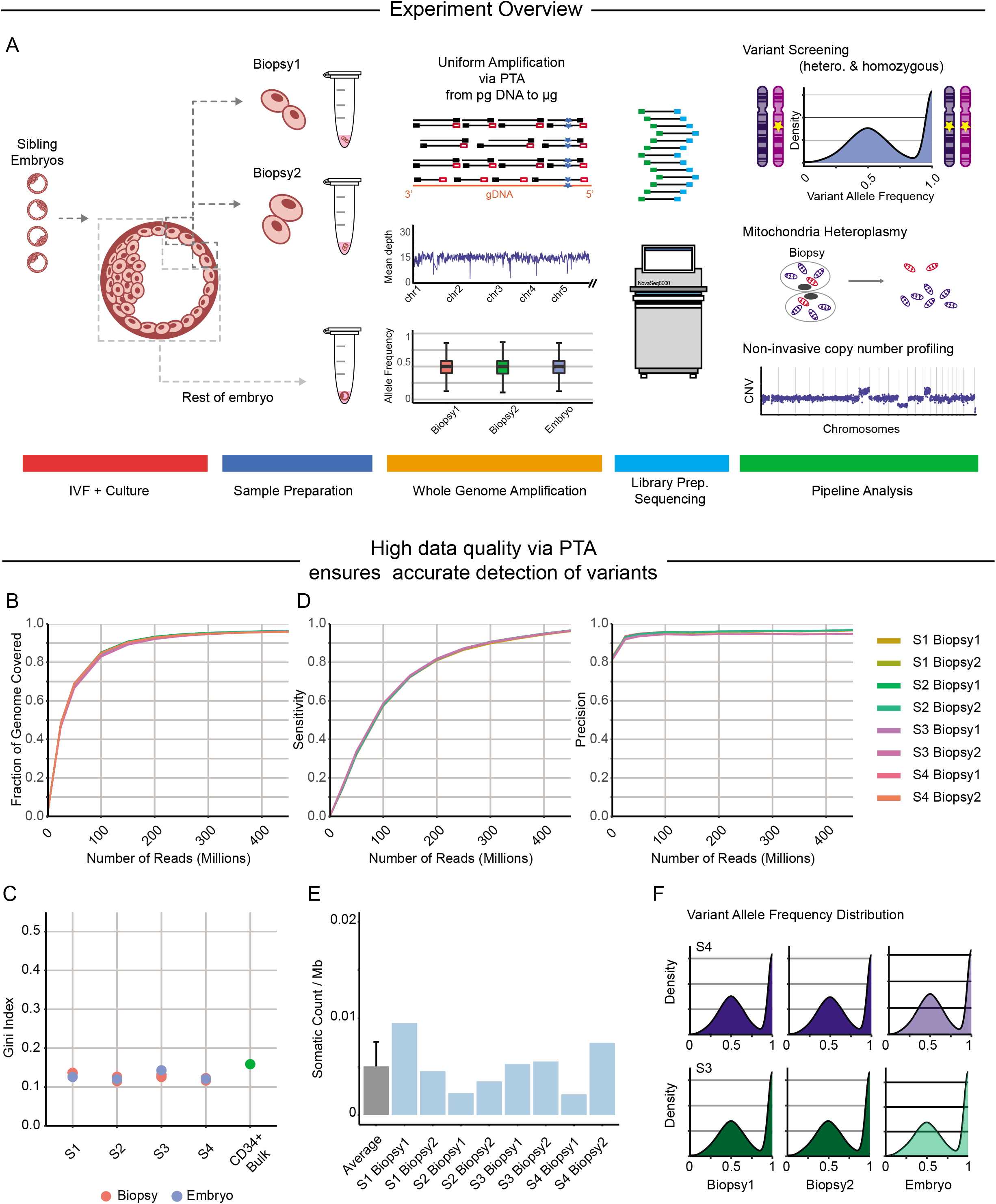
Experimental workflow and whole genome sequencing quality. A) A total of 4 sibling embryos were used in this study with 2 biopsies performed on each embryo. Standard PTA followed by sequencing were then performed on all biopsies. In parallel, the remaining embryos were processed to serve as a reference. Data analyses were performed to assess for aneuploidy, mendelian diseases, mitochondrial heteroplasmy, and polygenic risk scores in both the biopsies and spent media. B) There was approximately 96% genomic coverage in all biopsies at 450M reads (14X coverage). C) Gini Indices, which is used to assess amplification uniformity, were consistent between embryos and corresponding biopsies. The Gini index of amplified samples is comparable to an unamplified bulk sample. D) Sensitivity and precision of 96.3% and 96.2%, respectively, were observed among all samples at 450M reads. E) Somatic variants in biopsies are estimated to be 0.005 variants/Mb using SCAN2. F) Both heterozygous variants and homozygous variants were captured in biopsies, and the VAF distributions were similar across all sample types.

### Coverage, uniformity, variant calling, allelic balance, and reproducibility of the PTA-based PGT

Comparable genome coverage was achieved among the biopsies, embryos, and a bulk CD34+ cord blood sample when subsampled to the same read depth (Fig.1B & S1A). The coverage initially rises rapidly with increasing number of reads, followed by a coverage saturation at 96% with 450M reads, corresponding to a mean of 14X sequencing depth (Fig.1B). The uniformity was assessed by constructing Lorenz curve and associated Gini indices (GI) for each sample. Although biopsies of 5-8 cells contained 20 times fewer cells than the ~200 cell embryos, a mean GI of 0.13 was obtained for each biopsy, which was similar to the embryos, as well as the CD34+ cord blood bulk sample (Fig.1C & S1B). In each biopsy, we called an average of 3.27 million SNVs, among which 3.14 million were shared with the corresponding embryo (Fig.S1C). These correspond to an estimated sensitivity of 96.3% and precision of 96.2% (Fig.1D)^5^.

To estimate the somatic variants in each biopsy, we employed the somatic SNV caller SCAN2 that was recently developed for highly specific variant calling from samples that had undergone PTA.^6^ Using this approach we estimated a somatic variant burden of 0.005 variants/Mb, or about 15 variants per genome in the biopsies that were not detected in the corresponding embryo (Fig.1E). This suggests that the chromosomal instability seen in early embryos^7^ is not associated with the widespread acquisition of small genetic variants.

Allelic dropout due to loss of coverage or allelic imbalances, which are seen with previous WGA methods^5^, will have diminished sensitivity that could result in the loss of detection of pathological variants^3^. To assess this, we created variant allele frequency (VAF) histograms, which show that either biopsy has the same distribution as the corresponding embryo (Fig.1F & S2B). To verify the reproducibility of PTA in PGT, amplification of the 8 biopsies was conducted in three separate batches one week apart. As expected, we observed equivalent genomic coverage, uniformity, concordance, and allelic balance for all samples (Fig.1B-F).

### Detection of fetal aneuploidy

Our first analytic assessment was to determine if we could detect aneuploidy in the embryos, which is the most common use of PGT. As seen in Fig. 2, the two biopsies were concordant with the embryo in 3 of 4 cases. Interestingly, the one discordant case had one biopsy consistent with the embryo while the other showed a diploid profile. Upon closer examination of embryo S2 and the concordant biopsy, the loss of chromosome 14 was partial, suggesting one area of the embryo was mosaic for the loss of chromosome 14 while the area taken by the second biopsy was diploid, as has been previously reported (Fig.S2B)^8^.

**Figure 2.**
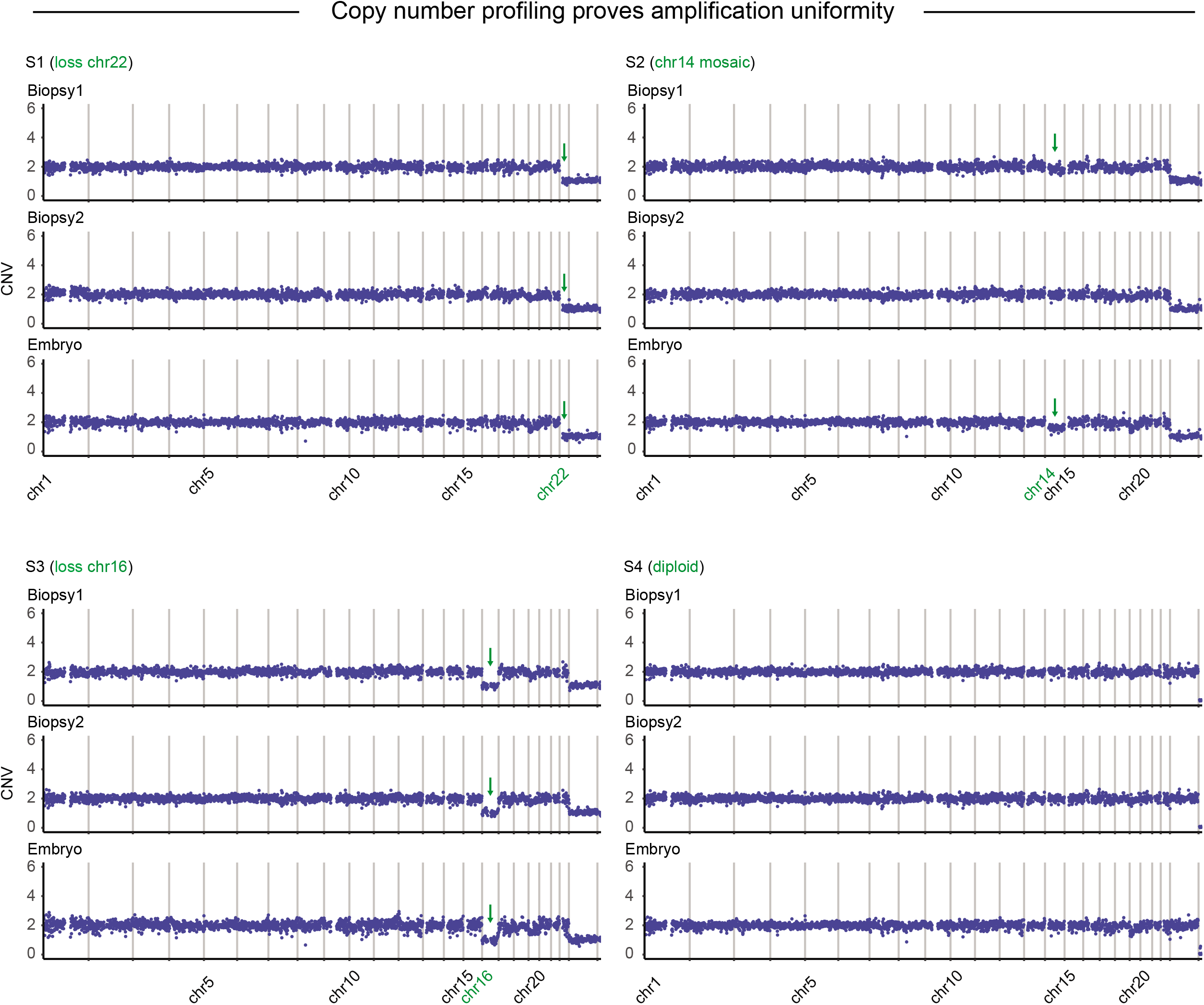
Copy number profiling. Aneuploidy screening was done using 1Mb bin size on 5M reads from each sample. Consistent copy number changes are observed with the exception of embryo S2 where mosaicism was detected at chromosome 14 in the embryo and one biopsy, but not in the remaining biopsy.

### Genome-wide screening for genetic diseases

We then sought to determine the additional genetic abnormalities that could be identified using our genome-wide approach. To achieve this, we primarily focused on variants that are rare in the population and reside in transcribed regions. Variants among those regions were then annotated for predicted consequences with MutationTaster^9^, and for known disease phenotypes using Clinvar^10^, HGMD^11^, and OMIM^12^. Importantly, the identified potential deleterious variants were concordant between both biopsies and the corresponding embryos (Fig.3A). Most of the disease-associated variants were also found in at least one additional embryo, suggesting they were inherited from one of the parents (Fig.3A). However, we also identified embryo-specific changes such as COQ8A A233T and ALOXE3 L237M, which may represent *de novo* variants.

**Figure 3.**
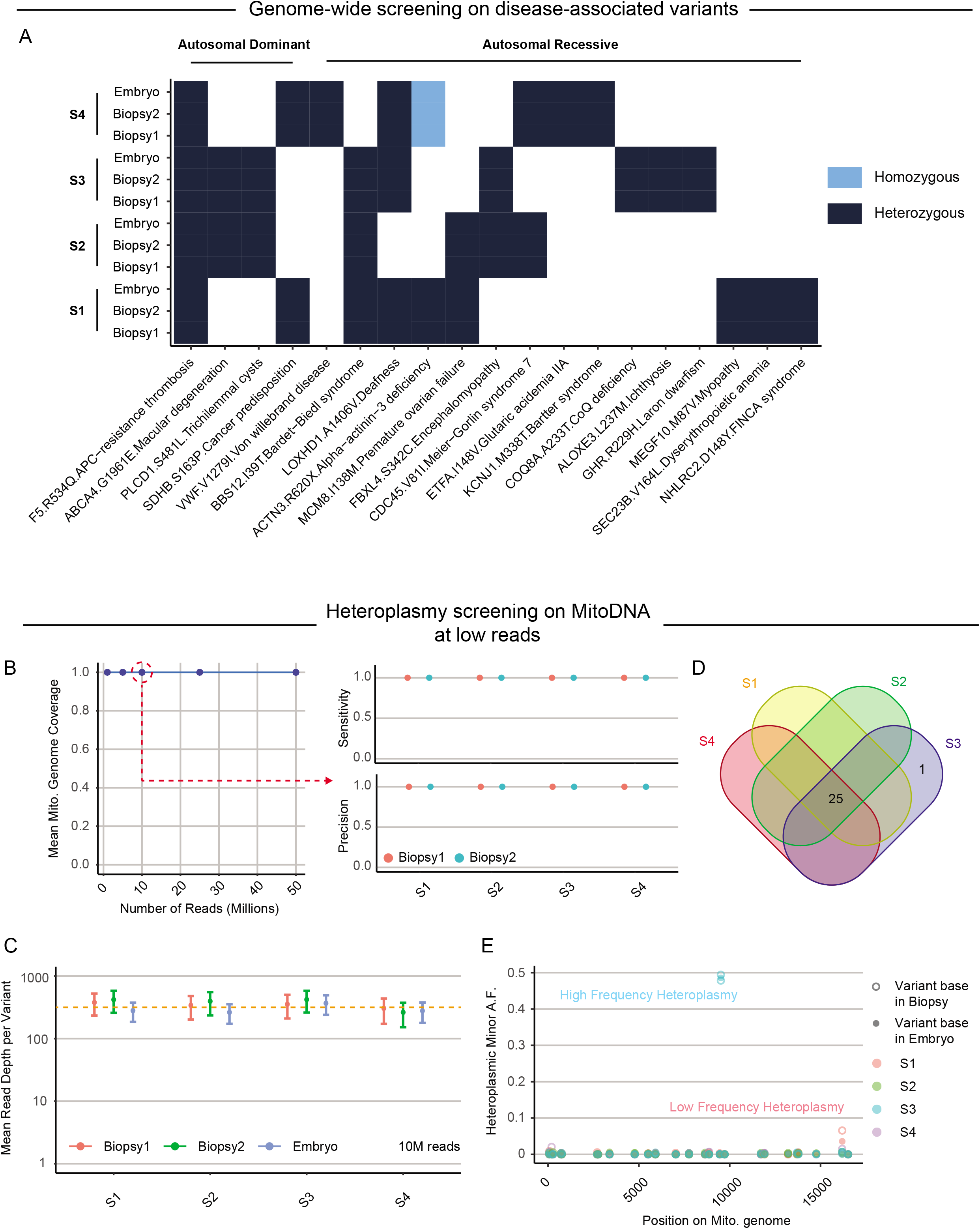
Genome-wide screening for disease-causing variants and mitochondria heteroplasmy. A) Genome-wide screening of known disease-associated variants. B) 100% of mitochondrial genomes were covered with 1M reads. Using 10M-read samples, sensitivity and precision reached 100% in both cases. C) A mean read depth of 342X was achieved with 10M reads. D) Venn diagrams indicate unique heteroplasmy in embryo of S3. E) Variants in biopsies and embryos were displayed as variant minor allele frequency across the mitochondrial genome.

We then focused on variants that have been described to cause autosomal dominant diseases (Fig.3A). This included a known gene conversion event in vWF (V1279I)^13^ that increases the risk of bleeding, SDHB S163P that has been shown to cause cancer predisposition^14^, and APC R534Q which increases risk for clotting^15, 16^. In addition, embryo S4 has autosomal recessive alpha-actinin-3 deficiency, which is associated with increased aerobic metabolism in skeletal muscle (Fig.3A)^17^. We also reported another 13 potential autosomal recessive variants here that are either known or adjacent to pathological variants (Fig.3A). Together, these data suggest it is feasible to screen almost the entire genome of an embryo for known disease-associated variants.

### Mitochondria heteroplasmic variant screening

Further analyses of the sequencing data from samples revealed 100% mitochondrial genome coverage with PTA with just 1 million reads, which is consistent with our previous work (Fig.3B)^5^. With 10 million reads, the mean sequencing depth of mtDNA was 342X, which is sufficient to detect heteroplasmy at approximately 1-2% frequency within each biopsy (Fig.3C). Meanwhile, both of sensitivity and precision reached 100%, indicating high concordance between mtDNA in embryos and biopsies using our approach (Fig.3C). As mtDNA are maternally inherited, we first confirmed high conservation of mtDNA among sibling embryos and their biopsies. All samples share the same 25 variants, including 5 in non-coding regions, 4 in rRNA regions, and 12 synonymous and 4 nonsynonymous coding variants (Fig.3D). None of these mtDNA variants have been reported as pathological^18^.

We then looked for embryo-specific mtDNA variants where we identified a unique heteroplasmic variant, C9512T in *COIII*, in embryo S3 (Fig.3D). It is synonymous and has not been reported to associate with known diseases. Importantly, this variant was identified in both biopsies and the embryo at similar frequencies (48-50%) (Fig.3E). It could have been inherited as *de novo* heteroplasmy in the egg or developed at a very early stage in the embryo, resulting in a selective advantage for those mitochondria. In addition, there is another low-frequency heteroplasmic variant in embryo S1 that is present at a mean of 4% allele frequency where the two biopsies possess 0% and 8% of this variant, respectively (Fig.3E). This spatial separation of that heteroplasmic variant between biopsies suggests it occurred in a mosaic population during the initial stages of embryogenesis. The copy number and mitochondrial variant calling data with just 10M reads suggests our approach allows us to accurately detect both heteroplasmy and aneuploidy with low-pass sequencing.

### Polygenic risk scores for 11 common diseases

Next, we directed our evaluation to common variants that are known to have small risk for common multifactorial diseases, and can be combined across the genome through the calculation of a polygenic risk score (PRS)^19^. With our broad genomic coverage, we successfully called more than 98% of selected SNP coordinates (7.5 million in total) from each biopsy, which is similar to the sensitivity of bulk PRS site coverage from 11 published studies (Fig.4A & S3A)^19–25^. Raw scores were then calculated and transformed into percentiles and prevalence using raw score distributions from the UK Biobank^26^, and whole embryos were processed in the same manner as controls. Importantly, we again saw consistent PRS percentile when comparing the embryo and corresponding biopsies in all cases, suggesting the feasibility of applying PRSs in PGT (Fig.4B-C & S3B-C).

**Figure 5.**
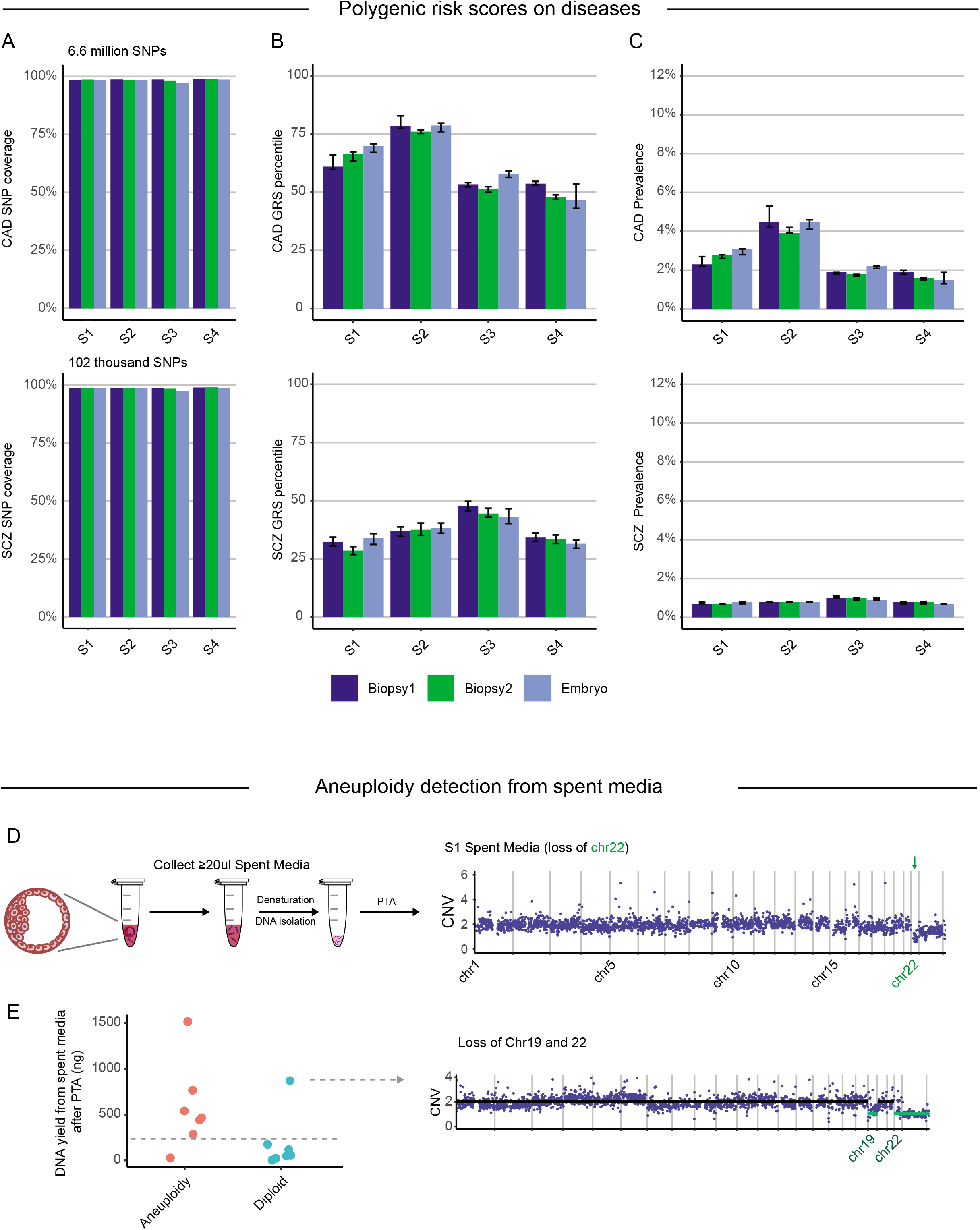
Polygenic risk scores for common diseases and Non-invasive aneuploidy assessment. A) 6.6 million and 0.1 million SNPs, respectively, were used to calculate PRSs for CAD and SCZ. We covered an average of 98.4% of SNPs for CAD and 98.6% for SCZ in each sample. B) Consistent PRS percentiles were seen between embryo and corresponding biopsies with percentiles varying between embryos C) The translation from percentile to prevalence reveals small absolute differences between embryos. D) We modified the PTA protocol to amplify DNA from spent media from the cultured embryos. CNV measurement from with spent media is consistent with CNV of embryos in aneuploid samples. E) DNA yield from each spent medium after PTA was plotted. Samples were classified into aneuploid and diploid based on each clinical PGT-A report of one biopsy. Spent media from aneuploid samples yielded about 3X more DNA. The outlier that was called diploid by clinical PGT-A had a loss of chromosomes 19 and 22 based on our analyses. Although further validation is needed, we see 86% concurrence with clinical PGT-A using only DNA yield after PTA.

Interestingly, coronary artery disease (CAD) and schizophrenia (SCZ) PRS-percentiles vary between embryos, even though they were derived from the same donors, which could be due to a combination of *de novo* variants in the embryo and the random co-segregation of risk alleles. (Fig.4B). In contrast, breast cancer and atrial fibrillation PRS-percentiles were generally low (Fig.S3A-C). We identified embryo S3 has an increased risk of type 1 diabetes (>=3X odd ratio). However, taking into account the low prevalence of type 1 diabetes in 50^th^ percentile controls (<0.9%), the probability of developing diabetes remains low^25^.

### Non-invasive aneuploidy assessment of spent media

It has been reported that aneuploidy can be detected noninvasively by amplifying DNA from spent media of embryos^27^. We hypothesized that using PTA we could not only detect aneuploidy but sequence a significant proportion of the embryo genome noninvasively. We therefore modified the PTA protocol to extract DNA from spent media prior to amplification (Fig.4D). As expected, CNV calling from the two available spent media samples (embryos S1 and S3) showed aneuploid profiles consistent with the biopsies (Fig.4D & S4A). Interestingly, we saw a trend of increased DNA yield from PTA in the aneuploid embryos when compared to diploid samples, suggesting that there is increased cell death in the aneuploid cells and that DNA yield alone could be sufficient for screening embryos. Additional trials were therefore performed and compared to the clinical reports where we again found that spent media of aneuploid embryos exhibited a roughly 3-fold increase in yield (Fig.4E). Interestingly, a diploid embryo based on clinical diagnostics (Fig4E-right) exhibited high DNA yield and loss of chromosomes 22 after sequencing using our approach, suggesting the original chromosomal ploidy estimation may have been inaccurate.

We then performed deep whole genome sequencing of the spend media where we found that genome coverage saturated at 50-70% with significant allelic drop out (Fig.S4B-C). In addition, we were unable to produce sequencable libraries from the diploid samples, which was likely the result of insufficient quantity of DNA in the media. Still, further work is needed to determine if there is sufficient DNA in the media of diploid embryos for noninvasive whole genome evaluations.

## Discussion

In this study, we outline a new strategy for genome-wide PGT with PTA where we can detect CNV, small genetic variants, and heteroplasmy. Most current clinical PGT testing evaluates embryos for aneuploidy or a very small number of selected genomic variants, but not both. This is due to the inherent limitations of currently used WGA methods that introduce method-specific artifacts, limiting the accuracy of downstream analyses for a given WGA method. Previous studies have performed whole genome sequencing of embryo biopsies using multiple displacement amplification (MDA) ^28, 29^. However, those studies were also hampered by the artifacts introduced by MDA, including loss of coverage and uneven coverage that hamper variant calling. Further, studies that utilize MDA typically screen an unknown number of amplified embryo biopsies using SNP arrays or other methods prior to selecting the top candidates for whole genome sequencing, making the clinical implementation of that approach impractical.

In the present study, we took advantage of the high coverage breadth and uniformity, as well as the low error rate and high reproducibility of PTA to perform accurate genome-wide variant calling of embryo biopsies. Importantly, we did not screen embryos prior to sequencing, enabling the potential immediate translation of this approach into clinical practice. In addition, we utilized the accurate genome-wide variant data to calculate polygenic risk scores for common diseases. Finally, we provided initial feasibility data for performing whole genome sequencing of spent media with PTA where we found aneuploid embryos have significantly higher levels of DNA, making the presence of high levels of extracellular DNA a potential marker for fetal aneuploidy.

There are a number of important limitations to our study. As with all whole genome sequencing studies, variant calling from the three billion locations in the human genome is not perfect. Further work is needed to balance the tradeoff between sensitivity and precision to provide optimal clinical insights from the data. Related to that concern, even with extraordinarily accurate variant calling, the interpretation of a given variant is frequently based on imperfect, and in many cases, no empirical supporting evidence. Those concerns are further amplified when using polygenic risk scores that use multiple associated genes that each have a small change in risk. Together, these observations highlight the computational framework that needs to be developed for the responsible clinical implementation of genome-wide PGT with PTA, as well as the importance of caution when interpreting the results.

The creation and future clinical utilization of whole genome sequencing of embryos also bring up a number of important ethical concerns. First, with all the challenges in the interpretation of the results, which variants should be reported back to the family? Should future parents only receive information on variants known to cause a specific disease that arises early in life, or should reports also include adult-onset diseases or even just an increased risk of those diseases? What if parents would like additional nondisease related insights from the data, such as the probability of having specific traits? All of these ethical challenges don’t take into account the quality of genomic variant annotation where there is significantly more information for those of European ancestry than all other populations, creating a disparity between people of different ancestries. Finally, there are important questions around parental consent: 1) Can parents provide consent with all the technical caveats in the interpretation? 2) If parents are more sophisticated in their understanding, can they consent to information beyond what the average person would understand? 3) What about the consent of the unborn child and any potential future consequences as a result of sequencing their whole genome? All of these challenges, and others, need careful consideration by the reproductive health community to create consensus around appropriate and ethical best practices.

In summary, we have presented a new strategy for genome-wide disease screening of embryos that is able to capture almost all the genetic variants in a sample with high precision. This approach is now technically feasible in a clinical setting, although numerous computational, interpretive, and ethical challenges remain to be addressed. Still, this approach provides a path for screening embryos for most genetic variants associated with diseases, with the potential to prevent the suffering caused by thousands of incurable genetic diseases.

## Materials and Methods

### Sample approval and collection

This study was approved by Stanford University IRB # 58757. Experiments were performed in accordance with protocol guidelines and regulations. The couple involved in this study had standard clinical PGT-A on each embryo, followed by a written consent to donate aneuploid embryos for research. An ethics consult was also conducted to discuss the study.

### Whole genome amplification through PTA

Embryos were first removed from tissue bank, followed by two biopsies from each embryo after thawing using standard clinical procedures. Biopsies and embryos were transferred into a 200ul PCR-tube containing 3ul of cell buffer (Bioskryb Genomics) before PTA. For media samples, DNA in the culture media (20ul-30ul in volume) was extracted with 1X AMPure beads (Beckman) with 5 minutes incubation followed by 80% ethanol wash twice. Then 3ul of EB buffer was added to elute the DNA from the beads before PTA. Beads were left inside the tube during PTA.

PTA was performed according to the manufacturer’s instructions (BioSkryb Genomics). The one modification was that after addition of lysis buffer, a 5-minute incubation on ice and another 5-minute incubation at room temperature were performed to maximize DNA denaturation. After PTA, the DNA was purified using 2X AMPure XP magnetic beads (Beckman). Yields were measured using the Qubit dsDNA HS Assay Kit with a Qubit 4 fluorometer according to the manufacturer’s instructions (ThermoFisher).

### Library preparation and sequencing

DNA sizes were first confirmed by running 1-2% Agarose E-Gel (Invitrogen). Then, 500ng of PTA product was used for library preparation with KAPA HyperPlus kit without the fragmentation steps. 2.5 μM of unique dual index adapter (Integrated DNA Technologies) were used in the ligation. 10 cycles of PCR were then used in the final amplification of biopsies and embryos, and 15 cycles were used for media samples if the PTA product was less than 500 ng. DNA concentration in the library was again quantified using the Qubit 4 dsDNA HS. Library sizes were confirmed through Agilent 4200 tapestation D1000 ScreenTape assay (Agilent Technologies). Sequencing runs were performed on a MiniSeq for QC and NovaSeq 6000 for WGS with 500ul of 1.6pM sample.

### Benchmarking Experiments Data Analyses

Sequencing data were trimmed using trimmomatic to remove adapter sequences and low-quality terminal bases, followed by GATK4 best practices with genome assembly of GRCh38. In Brief, fastq files were aligned with bwa mem, and then the corresponding bam files were processed with base quality score recalibration (BQSR) with duplicate marking before loading into HaplotypeCaller. The resulting vcf files were combined and genotyped with GATK4 combineGVCFs and GenotypeGVCFs, followed by variant quality score recalibration (VQSR). Annotation of variants was done by Annovar and HDMD. Quality metrics such, as genomic coverage, were acquired from the bam files after BQSR and MarkDuplicates using qualimap, as well as GATK AlignmentMetricsAummary and CollectWgsMetrics. Sensitivity and precision curves were generated using RTG Tools in R on the same VCF files. No regions were excluded from the analyses. The sensitivity and precision were calculated using RTG Tools by comparing each biopsy to its corresponding embryo. Default parameters were used for our analyses. Sensitivity was defined as number of variants shared between embryo and biopsy over total number of variants detected in that embryo. Precision was defined as number of variants shared between embryo and biopsy over total number of variants detected in that biopsy.

### Variant allele frequency histogram

Allele frequency was calculated using the number of ALT read divided by total read at that position. To increase the speed of the calculations, we randomly sampled 10000 variants to generate the histograms.

### Chromosomal copy number variation

Ginkgo was used for CNV calling using default parameters. All data were aligned to hg38/GRCh38 and samples were converted to bed files with bedtools prior to running with Ginkgo. A bin size of 1Mbp and independent segmentation were used for CNV calling.

### Disease screening

Disease screening was performed on variants mapped to exonic regions, including 5bp adjacent to stop/start codons, splicing regions and ncRNA regions. These variants were first extracted followed by annotation with Gnomad, Clinvar, HGMD, MutationTaster and OMIM. The criteria for identification included: 1) Gnomad AF_popmax < 10%, 2) positive Clinvar or HGMD annotations, and 3) MutationTaster without label of “polymorphism automatic” as these variants are known to be harmless. Afterwards, ~280 variants were left which were subjected to further analyses using data in Clinvar and HGMD. Pathological and variants with conflicting evidence were then reviewed manually.

### Polygenic risk score analysis

Embryos and biopsies were subjected to GATK HaplotypeCaller to generate gVCF files containing all coordinates on the genome including REF bases (via BP_resolution option). A total of 7.5 million unique genomic coordinates were focused on based on published PRSs for 11 diseases. Percent coverage in this case is defined as successfully extracted coordinates over total required coordinates for each disease. We calculated raw scores of 11 PRS and adjusted for 4 principal components of ancestry to minimize spurious ancestry associations in the resulting polygenic score. The percentile is reported based on the raw score distribution of about 100,000 UK Biobank participants. To investigate the uncertainty in the PRS percentiles due to missing coordinates, we first simulated random genotypes for missing SNPs and constructed 95% confidence intervals of PRSs for CAD and SCZ. Then, the mean±errs of PRSs were converted to percentile following the same principles. The prevalence was converted using published AUC values and PRS_to_Abs package, or extrapolated from prevalence vs percentile curve if AUC was not available.

## Code availability

Packages/software used in this study are all open-source. GATK4 from Broad institute is available at https://gatk.broadinstitute.org/hc/en-us. RTG tools from Real Time Genomics is available at https://www.realtimegenomics.com/products/rtg-tools. HGMD is provided by Stanford subscription via QIAGEN (a public version is available at http://www.hgmd.cf.ac.uk/ac/index.php). Annovar is available at https://annovar.openbioinformatics.org/en/latest. The whole genome sequencing analysis pipeline described in “Benchmarking Experiments Data Analysis” section is available on Gawad-lab github page https://github.com/Gawad-Lab/.

## Author Contributions

Y. Xia, V. Gonzales-Pena, B.R. Beher, and C. Gawad designed experiments. B.R. Behr, and P. Park contributed key materials, methods, and discussion. Y. Xia, D.J. Klein, J. Luquette and V. Reddy performed and analyzed experiments. L. Puzon and N. Siddiqui performed PRS analysis. Y. Xia, C. Gawad wrote the manuscript.

## Acknowledgements

We would like to acknowledge Stanford Center for Biomedical Ethics for their ethics consultation. C.G. is supported by Burroughs Wellcome Fund Career Award for Medical Scientists, an NIH Director’s New Innovator Award (1DP2CA239145), and the Chan-Zuckerberg Biohub.

---

**Figure S1.**
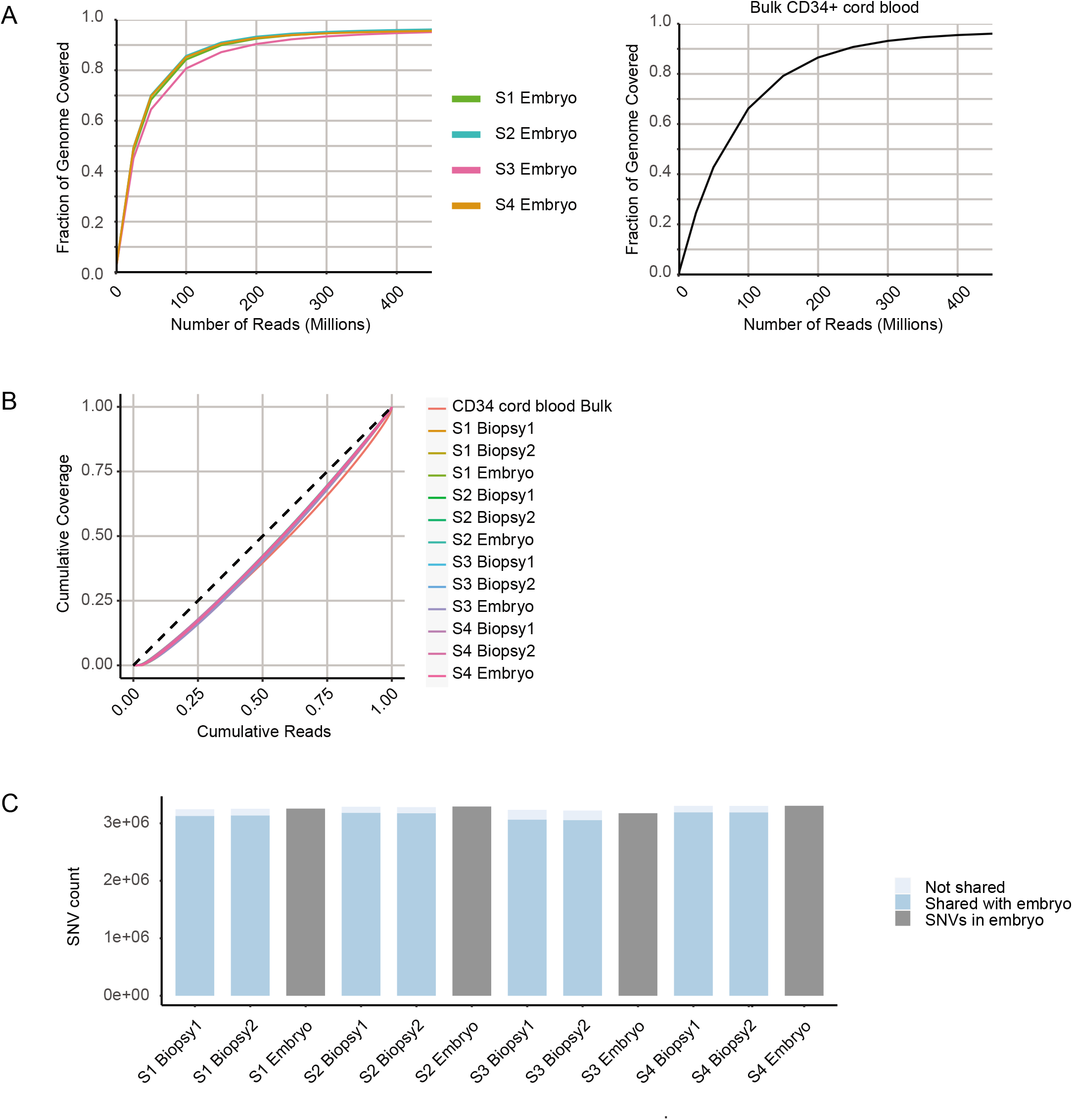
Additional embryo and biopsy genome coverage metrics. A) Genomic coverage of embryos and bulk CD34+ sample. B) Lorenz curves for all samples estimate the uniformity of amplification. C) An average of 3.27 million SNVs were called in each biopsy and 3.14 million of them were shared with the corresponding embryo.

**Figure S2.**
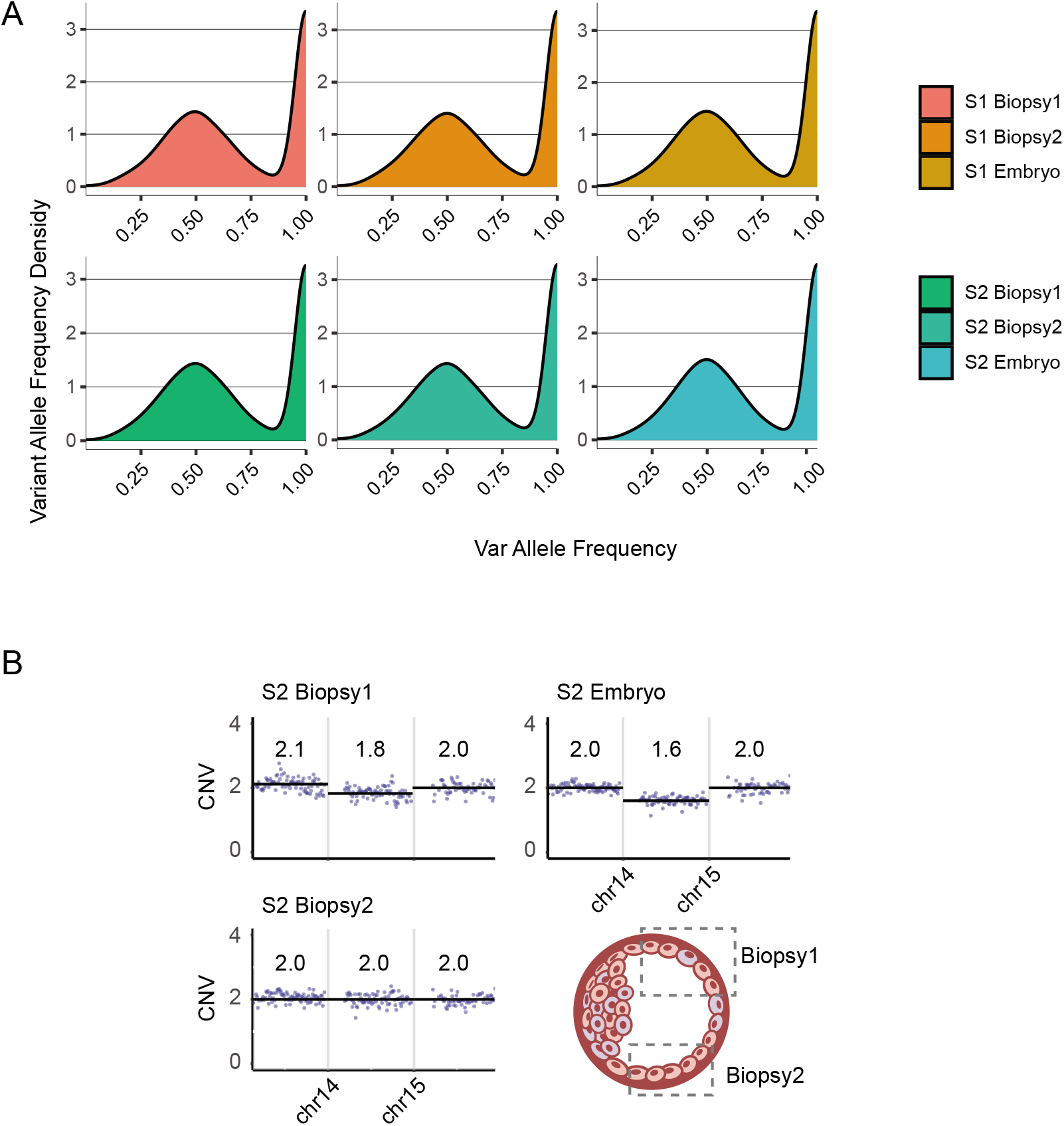
Allele balance and CNV in a mosaic embryo. A) Variant allele frequency is highly consistent among biopsies and corresponding embryos. B) An illustration of the CNV results in an embryo with mosaic loss of chromosome 14.

**Figure S3.**
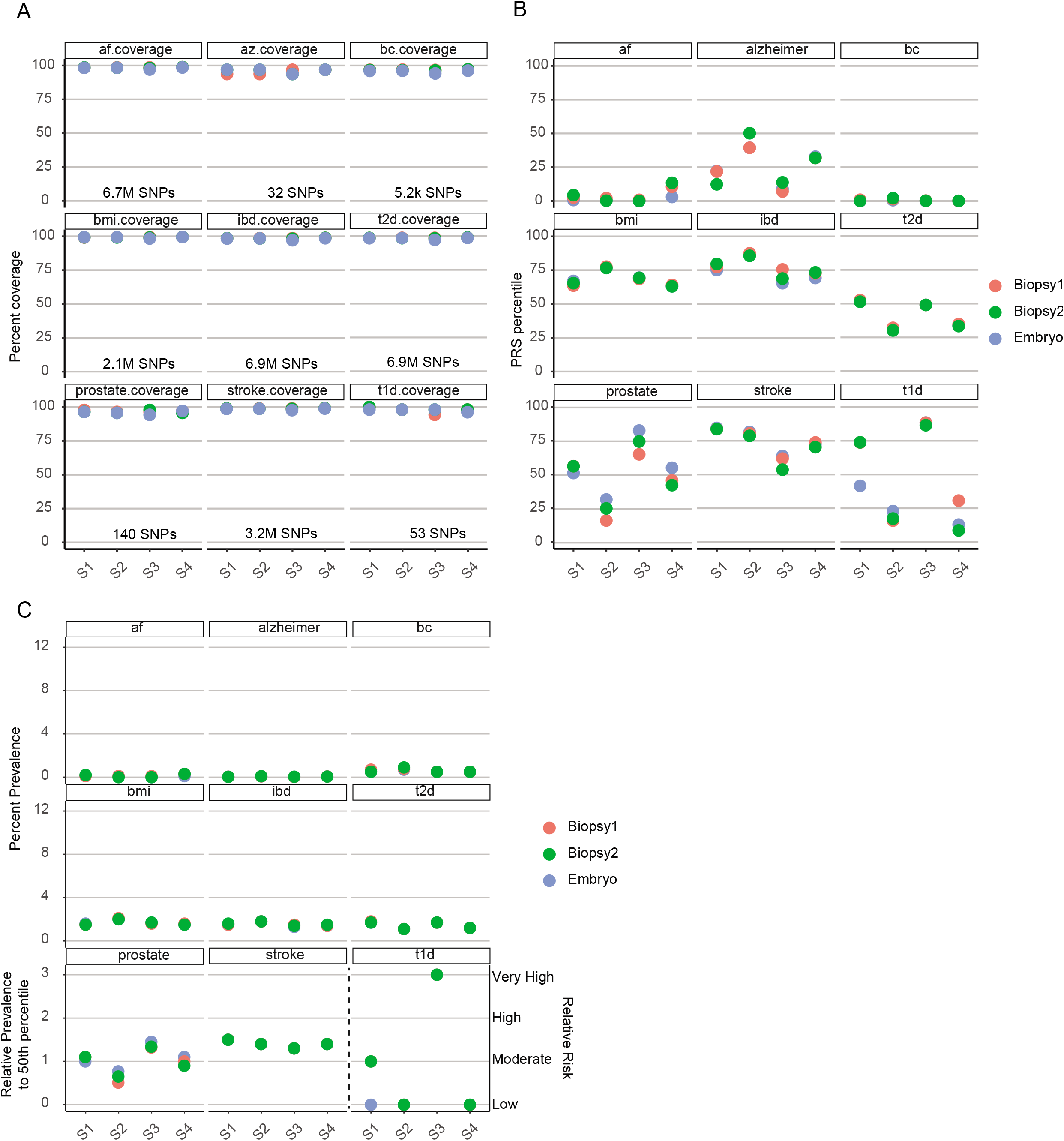
Polygenic risk score analyses of additional diseases. A) High target SNP coverage is observed for all 9 additional diseases. B) PRS percentile is plotted based on the polygenic risk sore of each sample. Consistent results between biopsies and corresponding embryos are observed. C) Percent prevalence or risk level is plotted based on the PRS percentile.

**Figure S4.**
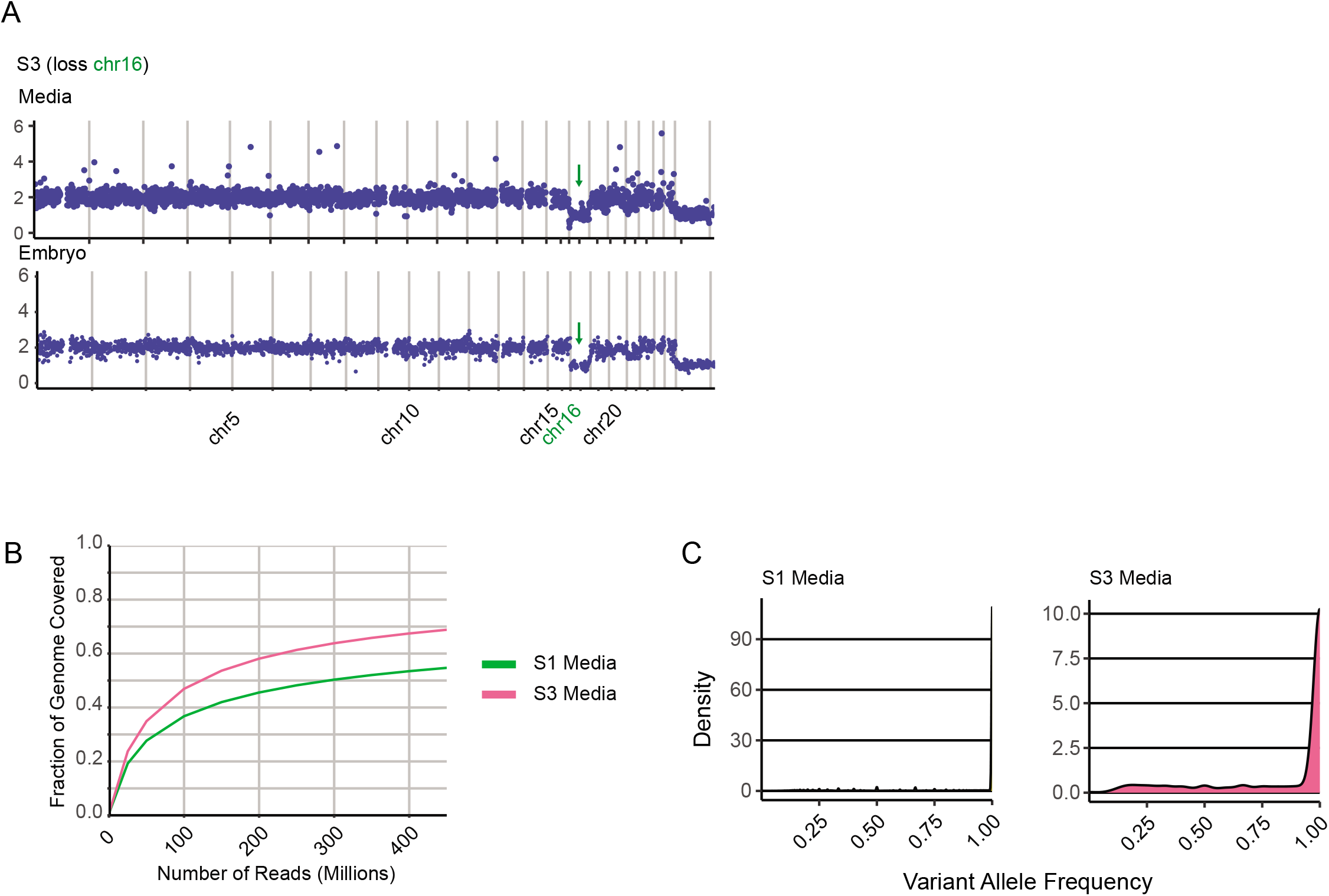
Non-invasive PGT-A. A) CNV from spent media is consistent with the CNV detected in each embryo. B) After amplifying DNA from spent media, partial genomic coverage is observed at 450M reads. C) One allele is almost universally lost in lower coverage sample S1, resulting in almost universal homozygous calls (VAF 1.0).

